# The length scale of multivalent interactions is evolutionarily conserved in fungal and vertebrate phase-separating proteins

**DOI:** 10.1101/2021.05.04.442641

**Authors:** Pouria Dasmeh, Roman Doronin, Andreas Wagner

## Abstract

One key feature of proteins that form liquid droplets by phase separation inside a cell is the presence of multiple sites – multivalency – that mediate interactions with other proteins. We know little about the variation of multivalency on evolutionary time scales. Here, we investigated the long-term evolution (∼600 million years) of multivalency in fungal mRNA decapping subunit 2 protein (Dcp2), and in the FET protein family. We found that multivalency varies substantially among the orthologs of these proteins. However, evolution has maintained the length scale at which sequence motifs that enable protein-protein interactions occur. That is, the total number of such motifs per hundred amino acids is higher and less variable than expected by neutral evolution. To help explain this evolutionary conservation, we developed a conformation classifier using machine-learning algorithms. This classifier demonstrates that disordered segments in Dcp2 and FET proteins tend to adopt compact conformations, which is necessary for phase separation. Thus, the evolutionary conservation we detected may help proteins preserve the ability to undergo phase separation. Altogether, our study reveals that the length scale of multivalent interactions is an evolutionarily conserved feature of two classes of phase-separating proteins in fungi and vertebrates.

## Body

Proteins that undergo liquid-liquid phase separation in a cell have various features that facilitate their condensation into liquid droplets. The presence of multiple interaction sites (multivalency) is one of these features. ^1-3^. Despite the pivotal role of multivalency, we know little about its evolution. The reason is that multivalency can take different forms, including the presence of interacting patches on a protein surface, short linear amino acid motifs, and specific amino acids within the intrinsically disordered regions of phase-separating proteins^4^.

We investigated the evolution of multivalency in two well-known classes of multivalent phase-separating proteins during ∼ 600 million years of evolution. The first class comprises orthologs of the fungal mRNA decapping subunit 2 protein (Dcp2). Dcp2 is one of the scaffold proteins that help RNA processing bodies (P-bodies) self-assemble by liquid-liquid phase separation^5-8^. P-bodies are conserved membrane-less eukaryotic organelles that contribute to the regulation of gene expression by participating in RNA decay and degradation^9^. They also serve as mRNA storage depots when cells are stressed^10^. Dcp2 undergoes multivalent interactions using short helical leucine-rich motifs (HLMs) in its disordered C-terminal domain^11^. HLMs form eight out of 12 identified interactions between Dcp2 and other core proteins in P-bodies^5^.

The second class of proteins comprises orthologs of six members of the FET family of RNA-binding proteins, including FUS, EWS, HNRNPA1, HNRNPA3, HNRNPR, and TAF15. These proteins have a common domain architecture that consists of a prion-like domain (PLD), and other domains with RNA/DNA binding affinities^10,12-14^. They contribute to DNA damage repair, transcriptional control, and the regulation of the life-time of RNAs in metazoan species^13^. The prion-like domain of these proteins has low sequence complexity, and is enriched in few amino acids, such as asparagine, glutamine, tyrosine, and glycine^15,16^. The aromatic residues within the prion-like domain, particularly tyrosine, and arginine in RNA binding domains, are responsible for the multivalency of these proteins^17-19^. Interactions between these residues drive the phase-separation of FET proteins^20^.

We use the stickers-and-spacers representation^21^ of phase-separating proteins throughout this work. Stickers are specific amino acids, motifs or protein domains whose interactions derive phase separation. Spacers are the sequences that separate the stickers. The helical leucine-rich motifs in Dcp2 and the aromatic residues in FET proteins are such stickers. For the FET proteins, we particularly focus on tyrosine residues, because their number and patterning in the sequence modulates the phase-separation propensity of these proteins^20,22,23^.

We first investigated the evolution of helical leucine-rich motifs in 48 Dcp2 proteins of the phylum *Ascomycota* (Dataset S1). HLMs lie within the disordered C-terminal domain of Dcp2 (Figure 1A, residues 229 - 930 in *S. cerevisiae*), and take the form LL-xϕ-L, where L stands for leucine, ϕ is a hydrophobic residue, and x represents any amino acid. We identified 347 motifs in these sequences that exactly matched the LL-xϕ-L pattern (Figure S1). As shown in Figure 1B, HLMs are the most conserved sequence segments within the intrinsically disordered C-terminal domain of Dcp2. However, their number substantially varies from a minimum of three in *L. elongisporus* to a maximum of 16 in *K. capsulata* (Table S1). Importantly, the number of HLMs increases with the length of the disordered C-terminal domain of a Dcp2 sequence (Figure 1C; Spearman correlation, R=0.44, p=0.0017). The average and median length of the spacer segments that separate HLMs are ∼70 and ∼51 amino acids. Based on these observations, we hypothesized that the scaling between the number of HLMs and the length of the disordered domain (1 HLM in ∼70 residues) reflects a requirement for a characteristic sequence length that separates sticker motifs. We tested this hypothesis by asking whether this characteristic sequence length may be subject to natural selection.

**Figure 1.**
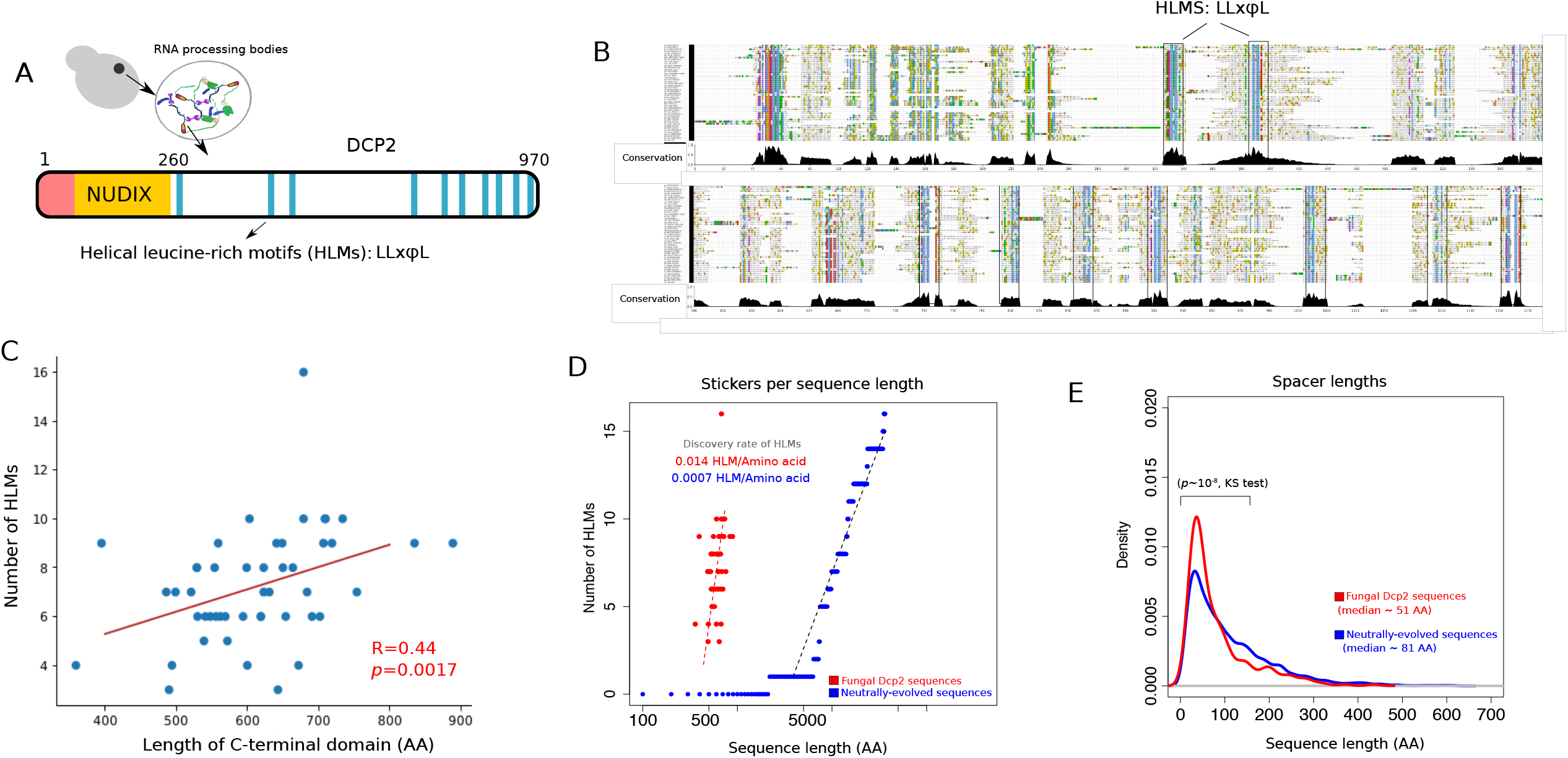
The length scale of multivalent interactions is evolutionary conserved in fungi species Dcp2. A) Architecture of Dcp2 in *S. cerevisiae* with the regulatory domain (in red), the NUDIX catalytic domain (in orange), and the disordered C-terminal domain (in white). Within the disordered C-terminal domain helical leucine-rich short linear motifs are responsible for the multivalency of Dcp2. B) Multiple sequence alignment of the C-terminal domain of Dcp2 in 48 fungal species within the phylum of *Ascomycota* spanning ∼ 600 million years of evolution. HLMs, shown as blue columns, are highly conserved within the C-terminal domain of Dcp2. C) The number of HLMs positively correlates with the length of the C-terminal domain of Dcp2 in fungi (Spearman correlation; R=0.44, p=0.0017). D) The incidence of HMLs in biological sequences (shown in red) is ∼ 35 times higher than that of neutrally evolved sequences (shown in blue). E) The distribution of spacer lengths (sequences that separate HLMs) in real Dcp2 sequences (shown in red), and in neutrally evolved sequences (shown in blue). We compared the two distributions and calculated the *p*-value for rejecting the null hypothesis that these distributions are indistinguishable by Kolmogorov-Smirnov (KS) test.

To understand the evolutionary forces that shape the scaling between the number of HLMs and the length of the C-terminal domain in Dcp2, we determined the likelihood that HLMs arise by chance through neutral evolution. To this end, we simulated neutral protein sequence evolution, using realistic divergence times of real Dcp2 sequences (see Method for details). We found that neutral evolution can indeed create motifs that exactly match known HLMs (Figure S2; Dataset S2), but the fraction of these neutrally-evolved HLMs per unit sequence length was much lower than that of HLMs in real sequences. Specifically, neutral evolution creates only one HLM per ∼1500 amino acids. In other words, HLMs in neutrally evolving sequences are ∼35 times less frequent than in real Dcp2 sequences (Figure 1D). We recalculated the fraction of HLMs per unit of sequence length for various codon frequencies, nonsynonymous substitution rates, and values of transition/transversion bias (Dataset S2). In all these calculations, we found a substantially higher incidence of HLMs per unit of sequence length in biological sequences compared to sequences evolved by neutral evolution (Table S2).

We also compared the distribution of spacer lengths (segments that separate HLMs) in the C-terminal domain of Dcp2 orthologs with that of neutrally evolved sequences. The median length of spacers is 81 amino acids in neutrally evolved sequences, significantly higher than the 51 amino acids in biological Dcp2 sequences (*p* ∼ 10^−6^; Wilcoxon ran-sum test; Figure 1E). In addition, spacer lengths are significantly more variable in neutrally evolved sequences compared to the biological Dcp2 proteins (*p* ∼ 10^−7^, one-sided F-test for the equality of variances), and the length distributions are significantly different (*p* ∼ 10^−8^; Kolmogorov-Smirnov test). Altogether, these results show that evolution has not only increased the incident of HLMs in Dcp2 sequences, but also has stabilized the lengths of sequences that separate HLMs.

To find out whether the scaling of sticker number with the length of a disordered region is a more general property, we next studied the FET family of proteins in vertebrates. We identified ∼200-300 orthologs for each of the six FET proteins, and compiled a set of 1480 sequences of these proteins (Dataset S3). Analogous to Dcp2 and its HLMs, we observed that longer FET proteins have more arginine (R) and tyrosine (Y) sticker residues in their prion like domain (Figure 2A, Spearman correlation, R=0.8, p<10^−16^). Importantly, among all 20 amino acids, the number of Rs and Ys showed the highest correlation with sequence length (Figure 2B; adjusted R^2^ ∼ 0.81, and 0.70 in a linear model with 5-fold cross-validation and 10 replicates). In sum, the scaling of multivalency with the lengths of disordered domains is not unique to Dcp2 in fungi. It also exists in the FET protein family of vertebrates.

**Figure 2.**
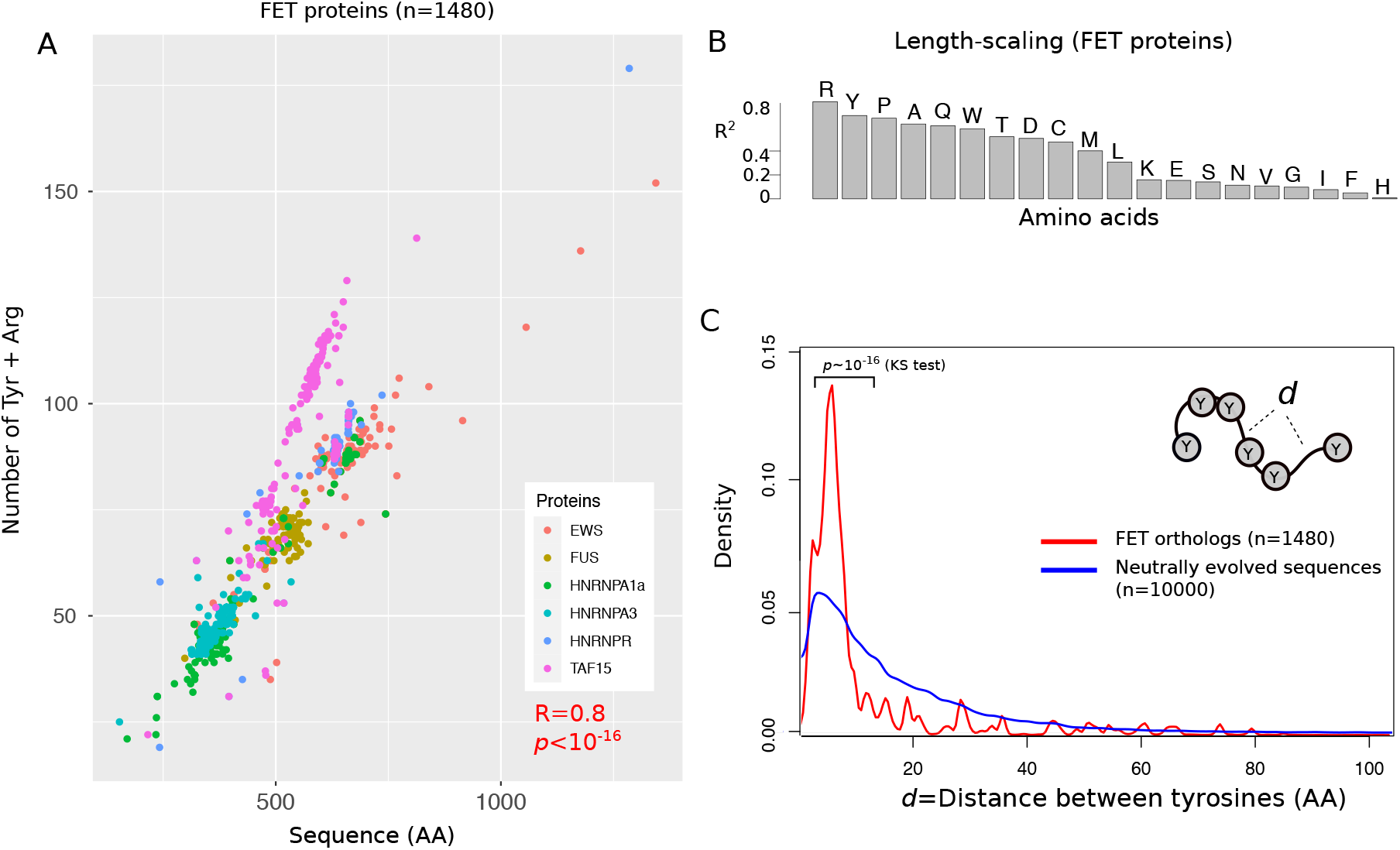
The length scale of multivalent interactions is evolutionary conserved in the FET family of vertebrate proteins. A) The number of arginine (R) and tyrosine (Y) residues of six different FET family members and their orthologs in vertebrate species (1180 proteins overall) versus their sequence length. B) The coefficient of determination (R^2^) between the number of different amino acids and the length of FET proteins and their orthologs. For a robust estimation of R^2^, we used a linear regression model with 5-fold cross-validation that we repeated 10 times. C) The distribution of distances between tyrosine residues in FET proteins (shown in red), and in neutrally-evolved sequences (shown in blue). We compared the two distributions and calculated the *p*-value for rejecting the null hypothesis that these distributions are indistinguishable by Kolmogorov-Smirnov (KS) test.

We further examined the spacer lengths that separate Ys and Rs in the sequence of FET proteins to find out whether natural selection has influenced the number of stickers per unit sequence length. We compared the distance distribution of both R and Y residues in FET proteins with that of neutrally-evolved sequences (see Methods for details). For both amino acids, the distribution of distances between tyrosine residues in FET proteins is significantly less variable than that of neutrally evolved sequences (See Figure 2C for spacers between tyrosine residues; *p* < 10^−16^; Kolmogorov-Smirnov test, and Figure S3 for spacers between arginine residues). The median distance between tyrosine residues is seven amino acids for FET proteins, which is significantly less than the corresponding distance of 11 amino acids in neutrally evolving proteins (*p* ∼ 10^−6^; Wilcoxon rank-sum test). In addition, the distance distribution of FET proteins is much more sharply peaked (leptokurtic, Figure 2C) and significantly differed from neutrally-evolved sequences (*p* ∼ 10^−8^; Kolmogorov-Smirnov test). This suggests that natural selection has likely stabilized this distance distribution in FET proteins.

Next we asked why the scaling of the number of stickers may be conserved, focusing on the hypothesis that it helps maintain a network of protein interactions that is necessary for condensation and phase separation^24^. To maintain this interaction network, disordered sequences should be able to adopt compact conformations. The reason is that only this type of conformation substantially increases the chance of interactions between stickers^24^. We thus wanted to find out whether this ability exists in our proteins. First, we calculated the fraction of charged residues in the spacers that separate HLMs in Dcp2 and aromatic residues in the prion-like domain of FET proteins. This fraction of charged residues is a proxy for the effective solvation and hence the conformation of disordered spacers^24^. Previous studies have suggested that spacers whose fraction of charged residues is less than 0.5 can self-associate and drive the formation of a condensation-promoting network of interactions (Figure 3A). As shown in Figures 3B-C, we found that almost all spacers in Dcp2 and FET proteins have a fraction of charged residues between 0.2 and 0.4, indicating that they can adopt compact conformations.

**Figure 3.**
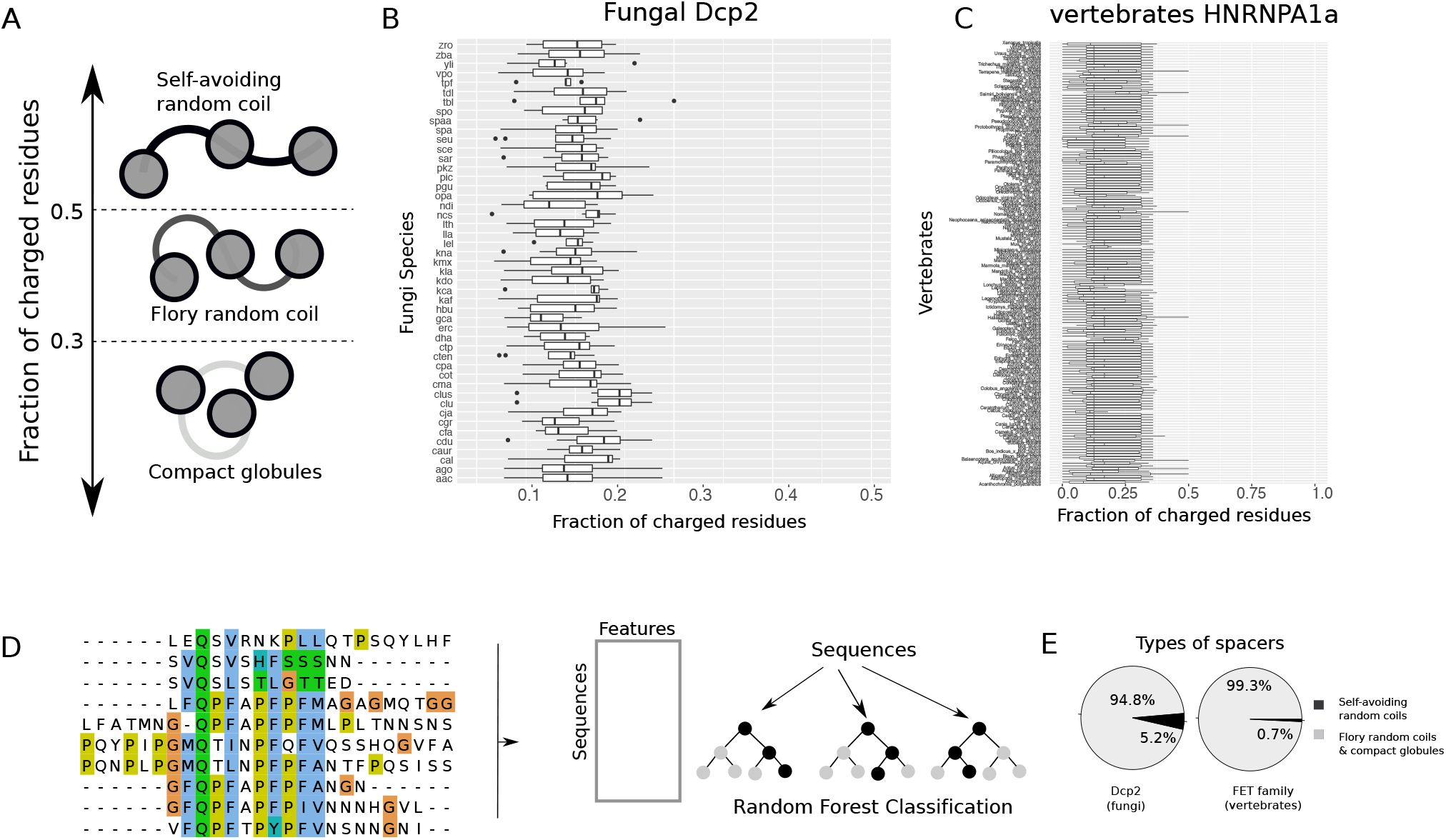
The disordered spacers in fungal Dcp2 and vertebrates FET proteins adopt conformations that promote phase-separation. A) The fraction of charged residues (FCR) can distinguish the conformation of spacer segments in multivalent proteins. Proteins with FCR > 0.5 preferentially adopt extended conformations like idealized self-avoiding random coils. Proteins with FCR < 0.3 can form compact globules. Sequences with intermediate values of FCR form conformations similar to Flory random coils where the net attractive and repulsive forces between residues and solvent molecules are in balance. The fraction of charged residues for B) fungal Dcp2 sequences, and C) vertebrate HNRNPA1a, a member of the FET family. D) Schematic for machine-learning random-forest classification to classify spacer types from their amino acid sequence. In brief, we used the sequences of naturally occurring disordered sequences that connect different domains, calculated the average of 500 amino acid properties for each sequence, and used this dataset to classify these sequences into the two categories of self-avoiding random coils, and Flory-random coils and compact globules. E) The fraction of spacers that adopt compact conformations (Flory random coils, and compact globules) and those that adopt extended conformations (self-avoiding random coils) in fungal Dcp2 and vertebrate FET proteins.

Second, we predicted a structural feature of disordered sequences known as the *Δ-*parameter. This parameter is the average difference between inter-residue distances of a disordered sequence and the corresponding distances of a typical Flory random coil^24^. Flory random coils are an idealized kind of disordered sequences in which the attractive and repulsive forces between residues and solvent molecules are at balance. Spacers that self-associate and promote phase-separation are characterized by *Δ* ≤ 0.1 nm. As *Δ* increases beyond 0.1 nm, spacers adopt more extended conformations, resembling another type of idealized sequence known as a self-avoiding random coil (Figure 3A).

We developed a sequence-based classifier of *Δ* using a random forest algorithm (Figure 3D), which classifies spacers based on their amino acid properties into two classes, those with *Δ > 0*.*1*, and those with *Δ* ≤ 0.1 nm (see Methods for details). We trained this classifier on a dataset of 256 naturally occurring disordered sequences whose *Δ* values had been previously calculated by molecular dynamics simulations^24^. This classifier achieved an accuracy of ∼ 0.88 in 100 independent runs with the data split into a training set (80% of the data) and a testing set (20%) (Figure S4 see Methods for details). Using this classifier, we found that ∼94.8% of all spacers in Dcp2 have a predicted value of *Δ* below 0.1 nm. We repeated this analysis for the spacers in the prion-like domain of the FET proteins and found that in these proteins too, most spacers (∼99.3%) have predicted *Δ* ≤ 0.1 nm. Altogether, these results indicate that both fungal Dcp2 sequences and vertebrate FET proteins have spacers that can self-associate and promote phase-separation in these proteins.

In summary, our work reveals that evolution has maintained a characteristic length scale of multivalent sticker sequences in two classes of multivalent proteins during ∼600 million years. Our results extend the previous observation by Martin *et al*. ^22^ that a uniform patterning of tyrosine residues in few members of FET proteins promotes phase-separation and inhibits the aggregation of these proteins. This scaling not only promotes phase-separation, but may also increase the robustness of proteins to DNA mutations such as indels and truncations. Dcp2 plays an important role in the assembly of RNA processing bodies, and FET proteins play such a role in the assembly of stress granules. Biomolecular condensates like these are sensitive to environmental stressors such as heat shock and energy depletion^16,25-27^. Our results thus also raise the intriguing possibility that evolution may have modulated the multivalency of proteins in membrane-less organelles to help organisms cope with new environments. To provide experimental support for this possibility is an exciting question for future work.

## Methods

### Data compilation and the generation of neutrally-evolved sequences

In this study, we used 48 orthologous coding sequences of fungal Dcp2, and ∼200-300 orthologs of six members of the FET family of proteins (FUS, EWS, HNRNPA1, HNRNPA3, HNRNPR, and TAF15; overall 1480 sequenced). We downloaded these sequences from the NCBI^28^, ENSEMBL^29^, and KEGG^30^ databases. Throughout, we worked with the amino acid sequences of these proteins, except for the simulation of neutral evolution, where we represented protein sequences on the level of DNA.

### Simulation of neutrally evolved sequences

We simulated protein evolution using the Evolver package within the PAML suite^31^. In brief, Evolver uses Monte Carlo simulations to generate codon sequences using a specified phylogenetic tree with given branch lengths, nucleotide frequencies, transition/transversion bias (*κ*), and the ratio of the rate of nonsynonymous to synonymous substitutions (d*N*/d*S*)^31^. To simulate neutral sequence evolution, we used the standard genetic code with codon frequencies from our study proteins sequences. Specifically, we used codon frequencies from the set of 48 Dcp2 sequences to model neutral evolution in fungal Dcp2, and codon frequencies from FUS orthologs for neutral evolution in vertebrate proteins. We used a consensus phylogenetic tree for the fungal species from the yeast genome browser^32^ and for the vertebrate species from the TimeTree database ^33^. To model neutral evolution, we set d*N*/d*S* to 1 and used a transition/transversion rate ratio of 2.3, and 2.9 for fungal and mammalian sequences. We estimated these values by fitting the codon model M1 to the phylogenetic tree and the sequences of these proteins. This model assumes that all branches of the phylogenetic tree have the same rate of evolution. We evaluated the number of neutrally-evolved HLMs for various values of d*N*/d*S* and the transition/transversion rate ratio to ensure that our results do not depend on the choice of these parameters (Table S1). Overall, we generated 10^4^ evolved sequences using this sequence evolution model.

### Detection of HLM motifs and their distinct flanking regions

We used regular expression matching to search for HLM motifs that matched the LL-xϕ-L pattern, where ϕ is a hydrophobic residue (one of the amino acids L, I, V, A, P, and F), and x represents any amino acid. To distinguish HLMs from HLM-like patterns we used the classification approaches of logistic regression and random forests implemented in the Python package scikit-learn. In these classifications, positive and the negative sets correspond to the flanking regions of HLMs and HLM-like motifs, respectively. The size of the training and the test set was 80%, and 20% of the whole dataset.

### Random forest classification of spacers

We used the random forest algorithm to develop a classifier of spacer conformation from the protein sequence. To this end, we used the average deviation of inter-residue distances of a spacer sequence from the same distances in a Flory Random Coil as the measure for the prediction of spacer types^24^. This deviation, known as the *Δ* parameter, can take positive and negative values. Disordered sequences with *Δ* ≤ 0.1 have the propensity to form compact conformations. We used a binary classification and classified proteins into a positive set (*Δ* ≤ 0.1) and a negative set (*Δ* > 0.1). To train our classifier we used a dataset of 256 disordered sequences for which we had calculated *Δ* by all-atom molecular dynamic simulations.

To build features for the classification, we calculated the average value of 500 physicochemical properties for each sequence in the positive and the negative sets. This yielded two feature matrices, one for sequences with *Δ* ≤ 0.1, and another for sequences with *Δ* > 0.1. To apply random forest classification, we used the randomForest package of R^34^, and evaluated the best number of trees (*nTree*) and the number of variables randomly sampled at each split (*mtry*) in the random forest algorithm. To do so, we systematically varied *nTree* and *mtry*, and calculated the accuracy of classification with 10-fold cross-validation in 3 replicates. We defined accuracy as the percentage of correctly identified classes of spacers (*Δ* ≤ 0.1 and *Δ* > 0.1) out of all spacers. The combination of *nTree*=5000 trees and *mtry*=10 variables achieved the highest accuracy of ∼ 88%. Here, we define accuracy as the ratio of the number of true positives to the sum of true positives and false negatives. We then used these parameters to perform 100 random forest clusterings, in which we randomly assigned proteins to the training and the testing datasets. To quantify the accuracy of classification we counted the number of true positive and false positive predictions and calculated the area under the curve (AUC). We represented these values by receiver operating characteristic curves (ROC) in Figure S2. We performed all statistical analyses using R. Scripts and input files for classification, as well as for evolutionary simulations are available at: https://github.com/dasmeh/multivalency_evolution

## Acknowledgement

The authors would like to thank Simon Alberti for careful reading of the manuscript and for helpful discussions on the evolution of liquid-liquid phase separation.

